# Assessment of DNA quality for whole genome library preparation

**DOI:** 10.1101/2024.03.20.585961

**Authors:** Linda Jansson, Siri Aili Fagerholm, Emelie Börkén, Arvid Hedén Gynnå, Maja Sidstedt, Christina Forsberg, Ricky Ansell, Johannes Hedman, Andreas Tillmar

## Abstract

In recent years, more sophisticated DNA technologies for genotyping have enabled considerable progress in various fields such as clinical genetics, archaeogenetics and forensic genetics. DNA samples previously rejected as too challenging to analyze due to low amounts of degraded DNA can now provide useful information. To increase the chances of success with the new methodologies, it is crucial to know the fragment size of the template DNA molecules, and whether the DNA in a sample is mostly single or double stranded. With this knowledge, an appropriate library preparation method can be chosen, and the DNA shearing parameters of the protocol can be adjusted to the DNA fragment size in the sample. In this study, we first developed and evaluated a user-friendly fluorometry-based protocol for estimation of DNA strandedness. We also evaluated different capillary electrophoresis methods for estimation of DNA fragmentation levels. Next, we applied the developed methodologies to a broad variety of DNA samples processed with different DNA extraction protocols. Our findings show that both the applied DNA extraction method and the sample type affect the DNA strandedness and fragmentation. The established protocols and the gained knowledge will be applicable for future sequencing-based high-density SNP genotyping in various fields.

## 1. Introduction

Efficient extraction of high-quality genomic DNA from limited sample amounts is a key challenge for cutting edge downstream applications like massively parallel sequencing (MPS). Numerous studies have evaluated and compared different DNA extraction methods [1-6], but the methodology for assessing DNA quality often relies on agarose gel electrophoresis and spectrophotometry which requires quite high amounts of DNA [7-9]. This may be problematic in fields such as forensic genetics, archaeogenetics, and clinical genetics, where the only samples available often contain low amounts of poor-quality DNA [10-15]. High-density single nucleotide polymorphism (SNP) genotyping through whole genome sequencing (WGS) or targeted hybridization capture sequencing have rapidly emerged as useful technologies in such fields [10-12, 16, 17]. While the use of microarrays has long been the state of the art for SNP genotyping, the requirement for high amounts of pure and intact DNA limits its application for several types of challenging samples [10, 18]. The recent progress in sequencing techniques has enabled the analysis of low quantities of degraded DNA [19-22] and the successful generation of genotypes from aged samples [1, 10, 14, 23-27]. The possibility of re-analyzing difficult DNA samples with more sophisticated methods has also spurred the continuation and reopening of unsolved investigations of “cold cases” and unidentified human remains [14, 24, 26]. However, low DNA quantity and quality often lead to reduced coverage, missing genotypes, and genotyping errors [14, 28]. The use of specific DNA extraction protocols may further augment DNA fragmentation [1, 2, 24, 27], possibly making an initial shearing step in whole genome library preparation protocols excessive [23]. Incomplete SNP results may also be due to having a large percentage of single-stranded DNA (ssDNA) in the extracts when applying a library preparation kit designed for double stranded DNA (dsDNA) [24, 29]. Attainable methodologies for assessment of DNA quality in terms of strandedness and fragmentation based on low DNA amounts are thus highly desired.

A major challenge when stored samples are re-analyzed with new technologies is that the DNA extraction methods initially applied were optimized for conventional approaches where DNA strandedness and fragmentation had minor impact on the outcome. For prospective sequencing-based genotyping, it is essential to know the proportion of dsDNA *versus* ssDNA and the level of DNA fragmentation prior to analysis. Only then, an appropriate library preparation kit optimized for either dsDNA or ssDNA can be chosen, input amount of DNA modified, and the DNA fragmentation parameters adjusted to the fragmentation level of the sample [30]. Thus, increased knowledge around the influence of relevant DNA extraction methods on DNA strandedness and fragmentation may greatly improve the success rate of downstream analyses.

We have developed, evaluated, and validated a set of analysis protocols for convenient determination of strandedness and fragmentation of DNA. Fluorometry measurements and linear regression modelling were applied to develop a protocol for estimation of the proportions of dsDNA *versus* ssDNA. Two different capillary electrophoresis methods were evaluated for determination of dsDNA fragment sizes and the performance of denaturing gel electrophoresis was applied for an approximate estimation of ssDNA fragment range. The developed methodology was used to determine the strandedness and fragmentation of a broad variety of both mock casework samples and forensic casework samples. Our aim was to determine the effect on DNA quality of various DNA extraction protocols and cell types and to highlight the importance of choosing an appropriate combination of methods for DNA extraction and library preparation. The developed methodology may be used prior to whole genome library preparation to optimize the chance of analytical success. In addition, it may be used to enable informed decisions on applicable DNA extraction methodology depending on downstream applications.

## 2. Materials and methods

### 2.1 Overview

The first scope of this study was to establish appropriate methods for estimation of DNA strandedness, *i.e.,* the proportion of dsDNA *versus* ssDNA, and DNA fragment size. For estimation of DNA strandedness, a protocol and a linear regression model based on fluorometry measurements of dsDNA and ssDNA with Qubit (Qiagen, Hilden, Germany) were developed and validated. For estimation of DNA fragment size, the performance of the capillary electrophoresis instruments Fragment Analyzer and TapeStation (Agilent Technologies, Santa Clara, CA, USA) were evaluated and compared to results from denaturing gel electrophoresis. Next, the established protocols were used to analyze the impact of different DNA extraction protocols and cell/sample types on DNA strandedness and fragment size. Human genomic DNA of high molecular weight (gDNA) and forensic mock casework samples (nasal secretion on cotton swabs, whole blood, saliva on adhesive tapes, bone, semen) were prepared and processed with various DNA extraction protocols. The fragment sizes provided by the Fragment Analyzer were compared to the degradation indices obtained from real-time quantitative polymerase chain reaction (qPCR) with PowerQuant System (Promega Corporation, Madison, WI, USA). After one year of storage in − 20 °C, all mock casework samples were re-analyzed applying the fluorometry-based model and Fragment Analyzer.

In addition to mock casework samples, 45 forensic casework samples from closed investigations were assessed for DNA strandedness and fragment size applying the established protocols. These samples were divergent in terms of age (collected 2001-2020), sample type (blood, semen, saliva, bone, non-visible traces), background (swab, paper, clothes, chewing gum etc.), and DNA extraction methods (phenol-chloroform protocols, Chelex-based protocols, PrepFiler Express) (see sample details in section 3.3).

This study was approved by the Swedish Ethical Review Authority (2023-02627-01) and was conducted in accordance with the guidelines of the Declaration of Helsinki.

### 2.2 Preparation of biological samples

Human genomic DNA of high molecular weight (Roche Diagnostics, Basel, Switzerland) were processed in a single replicate per DNA extraction protocol (Table 1). For preparation of mock casework samples, nasal secretion on cotton swabs, whole blood (with 1.8 mg K_2_EDTA/mL blood), saliva and semen were collected from anonymous volunteers with informed consent. Bone samples were collected from anonymized forensic autopsy cases. For each biological sample type, three replicates were prepared for DNA extraction according to Table 1. Selefa cotton swabs (OneMed, Malmö, Sweden) with nasal secretion were cut into microfuge tubes or Investigator Lyse&Spin Baskets (Qiagen) for DNA extraction on EZ1. Homogenized saliva (200 µL) was pipetted onto SceneSafe Fast adhesive tapes (SceneSafe, Burnham-on-Crouch, UK) which were left to dry for 1 hour prior to placing the tape with the saliva into microfuge tubes. Bone pieces were pulverized using IKA Tube Mill control (IKA-Werke GmbH, Staufen, Germany) and placed in 50 mL Falcon tubes. The prepared samples were processed according to Table 1 with the DNA extraction protocols described in section 2.3. For the initial step of assessing methods for estimation of DNA strandedness and fragment size, 50 µL saliva samples were prepared. Blood samples (10 µL) with and without 1.8 mg K_2_EDTA/mL blood were prepared for an experiment investigating the effect of EDTA on DNA quality.

**Table 1.**
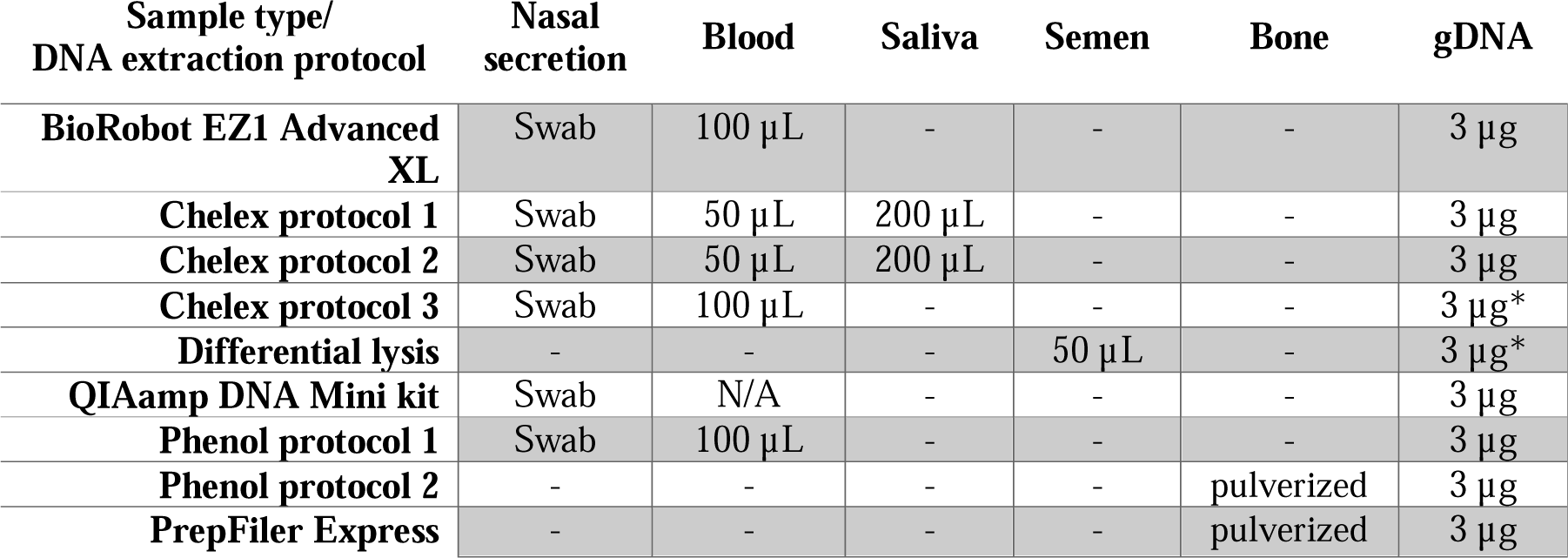
Overview of prepared mock casework samples and the applied DNA extraction protocols. All biological samples were prepared in triplicates and the gDNA in a single replicate per extraction protocol. Asterisk (*) indicates that the gDNA was added to the extraction tubes together with the Chelex solution, to avoid loss of DNA in the previous washing steps.

| Sample type/<br>DNA extraction protocol | Nasal<br>secretion | Blood | Saliva | Semen | Bone | gDNA |
| --- | --- | --- | --- | --- | --- | --- |
| **BioRobot EZ1 Advanced XL** | Swab | 100 µL | - | - | - | 3 µg |
| **Chelex protocol 1** | Swab | 50 µL | 200 µL | - | - | 3 µg |
| **Chelex protocol 2** | Swab | 50 µL | 200 µL | - | - | 3 µg |
| **Chelex protocol 3** | Swab | 100 µL | - | - | - | 3 µg* |
| **Differential lysis** | - | - | - | 50 µL | - | 3 µg* |
| **QIAamp DNA Mini kit** | Swab | N/A | - | - | - | 3 µg |
| **Phenol protocol 1** | Swab | 100 µL | - | - | - | 3 µg |
| **Phenol protocol 2** | - | - | - | - | pulverized | 3 µg |
| **PrepFiler Express** | - | - | - | - | pulverized | 3 µg |

### 2.3 DNA extraction and purification protocols

#### 2.3.1 BioRobot EZ1 Advanced XL

DNA extraction with EZ1 DNA Investigator kit (Qiagen) was performed according to the manufacturer’s instructions and the large volume protocol [31]. Briefly, 0.5 mL G2 buffer and 25 µL proteinase K was added to the samples prior to incubation on a thermoshaker at 56 °C, 900 rpm, for 1-18 hours. Samples were then centrifuged at 20,000 rcf for 5 min and the filter part with swab material removed. 400 µL of MTL buffer was added prior to placing samples in the EZ1 instrument. Elution volume was 50 µL.

#### 2.3.2 Chelex direct lysis protocol (Chelex protocol 1 and 2)

0.2 – 1 mL (sufficient to cover the sample) of 5% Chelex 100 Resin (Bio-Rad Laboratories, Hercules, CA, USA) with 0.2% Tween20 and 0.1 mg/mL proteinase K (Merck, Darmstadt, Germany) in Super-Q water was added to the microfuge tubes with samples [32]. For DNA extraction of mock casework samples, 800 µL was added to blood, 1 mL to samples with saliva on adhesive tape, 400 µL to the cotton swabs with nasal secretion, and 100 µL to the gDNA. Samples were incubated in ambient temperature for 30 min and briefly vortexed 3 times during this incubation. Mock samples were then incubated at 56 °C for either 45 (protocol 1) or 75 min (protocol 2), briefly vortexed, and then incubated for either 20 (protocol 1) or 40 min (protocol 2) at 100 °C in a heat cabinet. The forensic casework samples were incubated at 56 °C for 45 – 75 min followed by incubation at 100 °C for 20 – 40 min, according to a more flexible protocol. After incubation, the samples were allowed to cool down for 15 min at ambient temperature, before being briefly centrifuged at 11,000 rcf. Sample volumes of the adhesive tape samples were reduced to 200 µL using Amicon Ultra-2 30K tubes (Merck, section 2.3.9).

#### 2.3.3 Chelex protocol 3

1 mL of deionized water was added to the samples which were incubated in ambient temperature for 15-30 min and briefly vortexed 3 times during this incubation. Samples were then centrifuged at 11,000 rcf for 3 min and supernatant removed, leaving 30 – 50 µL in the tubes. 170 µL of 20% Chelex and 2 µL of proteinase K (10mg/mL) was added and samples were vortexed followed by two incubation steps at 56 °C for 75 min (mock casework samples) or 45 – 75 min (forensic casework samples) and 100 °C for 40 min (mock casework samples) or 20 – 40 min (forensic casework samples) with vortexing in between. After cooling down for 15 min at ambient temperature, samples were briefly centrifuged at 11,000 rcf.

#### 2.3.4 Differential lysis protocol

300 µL 0.5% digest buffer (10 mM Tris-HCl pH 7.5, 10 mM EDTA pH 8.5, 50 mM NaCl (Merck) and 0.5% sodium dodecyl sulfate, SDS (Invitrogen, Waltham, MA, USA)) and 60 µL proteinase K (10 mg/mL) were added to the samples. After a brief vortexing, samples were incubated at 56 °C for 30 min, then vortexed and centrifuged at 11,000 rcf for 5 min. 300 µL supernatant was removed (epithelial cell fraction) and 0.5 mL 2% digest buffer (10 mM Tris-HCl pH 7.5, 10 mM EDTA pH 8.5, 50 mM NaCl and 2% SDS) added to wash the remaining sperm fraction. After centrifugation at 11,000 rcf for 5 min, the supernatant was removed (leaving 30 – 50 µL) and the washing step with 2% digest buffer was repeated twice. 1 mL deionized water was added, and samples were centrifuged at 11,000 rcf for 5 min and supernatant removed (leaving 30 – 50 µL). 170 µL of 5% Chelex, 8 µL proteinase K (10 mg/mL) and 8 µL of 1 M DTT (Merck) was added to the sperm fraction which were vortexed and incubated at 56 °C for 60 min (mock casework samples) or 30 – 60 min (forensic casework samples). Samples were vortexed and incubated at 100 °C for 40 min (mock casework samples) or 20 – 40 (forensic casework samples), then vortexed again and briefly centrifuged at 11,000 rcf. Samples were then purified using Amicon Ultra-2 30K tubes (section 2.3.9).

#### 2.3.5 QIAamp DNA Mini kit

DNA extraction was performed according to the manufacturer’s protocol [33], with 200 µL elution volume. The swabs were removed after heat incubation at 56 °C for 2 hours by transferring samples including swab material to QIAshredder tubes (Qiagen) and centrifugation at 12,000 rcf for 2 min.

#### 2.3.6 Phenol protocol 1

0.5 mL 2% digest buffer and 15 µL proteinase K (10 mg/mL) was added to the samples. Samples were vortexed and incubated at 56 °C for 16 hours (mock samples) or 6 – 24 hours (forensic casework samples). Samples were briefly vortexed and centrifuged at 11,000 rcf, then transferred to a Phase Lock Gel (PLG) tube (QuantaBio, Beverly, MA, USA) containing 0.5 mL phenol:chloroform:isoamyl alcohol (25:24:1) buffer (Merck). After thorough vortexing, samples were centrifuged at 11,000 rcf for 5 min, and additional 0.5 mL phenol:chloroform:isoamyl alcohol buffer added. Thorough vortexing and centrifugation at 11,000 rcf for 5 min was repeated and the upper water phase containing DNA was carefully transferred to a microfuge tube with 0.5 mL H_2_0 saturated n-butanol (Merck). Tubes were vortexed and centrifuged at 11,000 rcf for 5 min, and the lower DNA containing phase was transferred to Amicon Ultra-2 30K tubes for purification (section 2.3.9).

#### 2.3.7 Phenol protocol 2

6 mL bone extraction buffer (1 M Trizma base, 0.1 M NaCl, 50 mM TitriplexIII (Merck), 0.5% SDS (Life Technologies, Carlsbad, CA, USA)) and 150 µL proteinase K (2 mg/mL) was added to the samples with pulverized bone. Tubes were shaken, incubated at 56 °C for 2 hours, then centrifuged at 2,000 rcf for 15 min. The upper phase was carefully poured into 15 mL MaXtract high density gel tubes (Qiagen) and 4 mL of phenol:chloroform:isoamyl alcohol (25:24:1) buffer added. Tubes were vigorously shaken and centrifuged at 1,500 rcf for 5 min. 4 mL of phenol:chloroform:isoamyl alcohol buffer was added, and tubes vigorously shaken and centrifuged at 1,500 rcf for 5 min. The upper phase was poured into an Amicon Ultra-2 30K filter device which were centrifuged at 3,500 rcf for 10 min. Flow-through was discarded and 3 mL Milli-Q water was added, the tubes were centrifuged at 3,500 rcf for 10 min. Samples were transferred to QIAquick tubes for purification according to QIAquick PCR purification kit protocol (Qiagen). Purification was performed according to the manufacturer’s instructions [34], with the modification that two purification steps with 650 µL PE buffer was performed prior to elution in 200 µL.

#### 2.3.8 PrepFiler Express BTA Forensic DNA extraction kit

DNA extraction was performed according to the bone and tooth protocol accompanying PrepFiler Express BTA Forensic DNA extraction kit (Applied Biosystems, Life Technologies Corporation, Carlsbad, CA, USA) [35]. Elution volume was 200 µL.

#### 2.3.9 Purification and volume reduction with Amicon Ultra-2 30K filter devices

Amicon Ultra-2 30K filter devices were used to reduce the volumes of adhesive tape samples and to purify the samples processed with differential lysis and phenol protocol 1. Sample extracts were added to the Amicon tubes together with low TE buffer (10 mM Tris-HCl, 0.1 mM EDTA, pH 8.0, Medicago, Uppsala, Sweden) to a total volume of 2 mL. Amicon tubes were centrifuged at 4,000 rcf for 15 min, flow-through discarded and additional 2 mL low TE buffer added to the filter tubes. After centrifugation at 4,000 rcf for 15 min, flow-through was discarded and filter devices inverted and centrifuged at 1,000 rcf for 2 min. Low TE buffer was applied to adjust the final sample volumes to 200 µL.

#### 2.3.10 Preparation of DNA dilutions

Following DNA extraction, the samples were quantified by qPCR (RB1 assay [36], details in section 2.4) and dilutions with concentrations of 0.5, 2 and 5 ng/µL DNA were generated for each extract. For blood samples extracted with Chelex protocol 1 and 2, DNA dilutions of 5 ng/µL could not be generated due to low DNA yields for this combination. The highest DNA concentration for these Chelex-processed blood samples were thus 2 ng/µL. Extraction blanks (DNA extraction performed without any biological material) were used to dilute the samples. This was done to preserve any effects on the measurements from the DNA extraction background. All extracted samples and dilutions were stored at − 20 °C until further analysis.

### 2.4 DNA quantification with qPCR

DNA quantification of the mock casework extracts was performed on a CFX96 Touch real-time PCR instrument (Bio-Rad Laboratories). The RB1 qPCR assay [36] was applied with 10X Immobuffer (Meridian Bioscience, Cincinnati, OH, USA), 0.2 mM dNTP (Roche Diagnostics), 4 mM MgCl2 (Roche Diagnostics), 0.3 µM of each primer (RB1_80F and RB1_235R, 0.2 µM hydrolysis probe (Integrated DNA Technologies, Coralville, IA, USA), 2 µg BSA (Roche Diagnostics) and 1 U Immolase DNA polymerase (Meridian Bioscience) in a total volume of 20 µL (2 µL sample). The qPCR protocol included DNA polymerase activation at 95 °C for 10 min, followed by 45 cycles at 95 °C for 10 s, 60 °C for 20 s and 72 °C for 30 s.

DNA quantification of forensic casework samples was performed applying PowerQuant System [37] on the real-time PCR instrument QuantStudio 5 (Thermo Fisher Scientific, Waltham, MA, USA). DNA quantification was based on amplification of the smaller autosomal target (84 bp) and a degradation index (DI) was generated from the ratio [small target] / [large target (294 bp)]. The mock casework samples of different DNA concentrations (0.5, 2 and 5 ng/µL) were also analyzed applying PowerQuant System, to obtain DI values. Standard curves based on 1:10 dilutions of 2800 M control DNA (Promega Corporation) (0.001 – 10 ng/µL, limit of quantification (LOQ) 0.001 ng/µL) were included for both DNA quantification protocols.

### 2.5 Determination of DNA strandedness and DNA fragmentation

#### 2.5.1 Measurements of dsDNA and ssDNA using Qubit

A protocol based on fluorometry was developed to estimate the proportion of dsDNA *versus* ssDNA in a DNA extract. The ratio of DNA concentrations as measured with Qubit dsDNA assay kit and Qubit ssDNA assay kit (Thermo Fisher Scientific) was assessed. Since the fluorescent dye of the Qubit ssDNA Assay Kit binds not only to ssDNA but also to dsDNA, the dsDNA was removed by treating the samples with dsDNAse prior to measuring ssDNA. First, the DNA extracts (2 µL subsample) were quantified using Qubit dsDNA HS assay kit according to the manufacturer’s manual [38]. Prior to quantification with Qubit ssDNA assay kit, 2 µL subsamples were treated with shrimp dsDNase (Thermo Fisher Scientific) according to protocol (incubation at 37 °C for 5 min) to remove all dsDNA [39]. The total volume (10 µL) of the dsDNase-treated sample was added for measurement with Qubit ssDNA assay kit [40]. Measurements were performed 9-11 min after the sample extract was added to the Qubit working solution to acquire stable values with both Qubit assays (ssDNA and dsDNA). A Qubit 3.0 instrument was used for all mock casework samples, and a Qubit 2.0 instrument was used for the forensic casework samples.

#### 2.5.2 Model generation for estimation of DNA strandedness

A linear regression model for estimation of the proportion of dsDNA in a sample as a function of the ratio of the DNA concentration given by Qubit dsDNA HS assay kit and the concentration given by Qubit ssDNA assay kit (after treatment with dsDNase) was established based on reference samples with known proportions of dsDNA and ssDNA. Human gDNA of high molecular weight (Roche Diagnostics) and ssDNA (M13mp18 single-stranded virion DNA, TaKaRa Bio, Kusatsu, Japan) were added in proportions of (in %) 0:100, 5:95, 10:90, 15:85, 20:80, 25:75, 30:70, 35:65, 40:60, 45:55, 50:50, 55:45, 60:40, 65:35, 70:30, 75:25, 80:20, 85:15, 90:10, 95:5, and 100:0 to generate samples with total DNA concentrations of 0.5, 2 and 5 ng/µL (n = 63). The model was validated based on mean absolute error (MAE) with a second set of reference samples, with dsDNA and ssDNA mixed in proportions of (in %) 0:100, 5:95, 10:90, 25:75, 50:50, 75:25, 90:10, 95:5, and 100:0 with total DNA concentrations of 0.5, 2 and 5 ng/µL (n = 26). By analyzing dilutions of saliva samples extracted with QIAamp DNA Mini kit and Chelex protocol 2 (two-fold serial dilutions from 0.0095 - 5 ng/µL as quantified by qPCR (RB1 assay, n = 3 per protocol and dilution), the LOQ was determined along with the precision of the method, assessed based on the coefficient of variation (CV).

#### 2.5.3 Assessment of DNA fragmentation

To assess DNA fragmentation, samples were analyzed on Fragment Analyzer with Genomic DNA 50kb Kit and TapeStation with either High Sensitivity D5000 ScreenTape System or High Sensitivity RNA ScreenTape System (Agilent Technologies) according to the manufacturer’s protocols [41-43]. Average fragment size, peak width, and percentage of DNA in the two major peaks along with the genomic quality number (GQN) with a fragment size threshold of 10 kb were obtained by Fragment Analyzer. Additionally, extracts with high percentages of ssDNA were analyzed on denaturing gels (1 M urea, 0.8% agarose, 1x Tris-acetate-EDTA (TAE) buffer and 1x SybrGold (Thermo Fisher Scientific)) at 50 V for 3 hours. To resolve any secondary structures, the samples were mixed with NorthernMax formaldehyde-based loading dye (Thermo Fisher Scientific) and heated to 65 °C for 15 min, prior to loading the gel.

The LOQ of Fragment Analyzer (50kb kit) and TapeStation (D5000 kit) were determined by analyzing dilutions of saliva extracts processed with QIAamp DNA Mini kit (two-fold serial dilutions from 0.0095 - 5 ng/µL as quantified by qPCR, n = 3 per dilution). The precision (CV) of DNA fragment size data generated by Fragment Analyzer was also determined. DI values based on the PowerQuant amplification of two PCR targets of different lengths were generated for both mock and forensic casework samples.

#### 2.5.3 Statistical analyses

The mean absolute error (MAE) was determined by

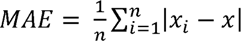

where n = number of measurements and |*x_i_ - x*| = the absolute error (absolute difference between true and estimated value). One-way ANOVA followed by Bonferroni post hoc tests (*i.e*. significance level 0.05 divided by the number of comparisons), was applied to investigate whether dilutions of DNA extracts processed by Chelex protocol 2 and QIAamp gave equivalent [dsDNA]/[ssDNA], average fragment sizes and GQN, respectively. The data sets subjected to one-way ANOVA met the criteria of approximately normal distributed data (evaluated by inspection of residuals histograms) and approximately equal variances (tested by applying Levene’s test in R package car [44, 45]. Heteroscedastic two-tailed t-tests were applied for 1) comparison of average DNA fragment sizes and GQN in saliva samples processed with QIAamp DNA Mini kit and Chelex protocol 2, 2) investigations of differences in [dsDNA]/[ssDNA], fragment size or DI:s between mock and forensic casework extracts from the same sample types, and 3) differences in DI values between forensic casework samples derived from non-visible traces and blood samples. A power regression analysis was performed to describe the relationship between average DNA fragment sizes and the DI values. Two-tailed paired t-tests with a significance level of 0.05 were applied to assess whether the percentage of dsDNA and average fragment sizes had changed after one-year storage in – 20 °C.

## 3. Results and discussion

### 3.1 Development and evaluation of methods for estimation of DNA quality

The first scope of this study was to develop and establish useful methods for estimation of DNA quality prior to whole genome library preparation, overview of protocols and workflow in Fig 1.

**Fig 1.**
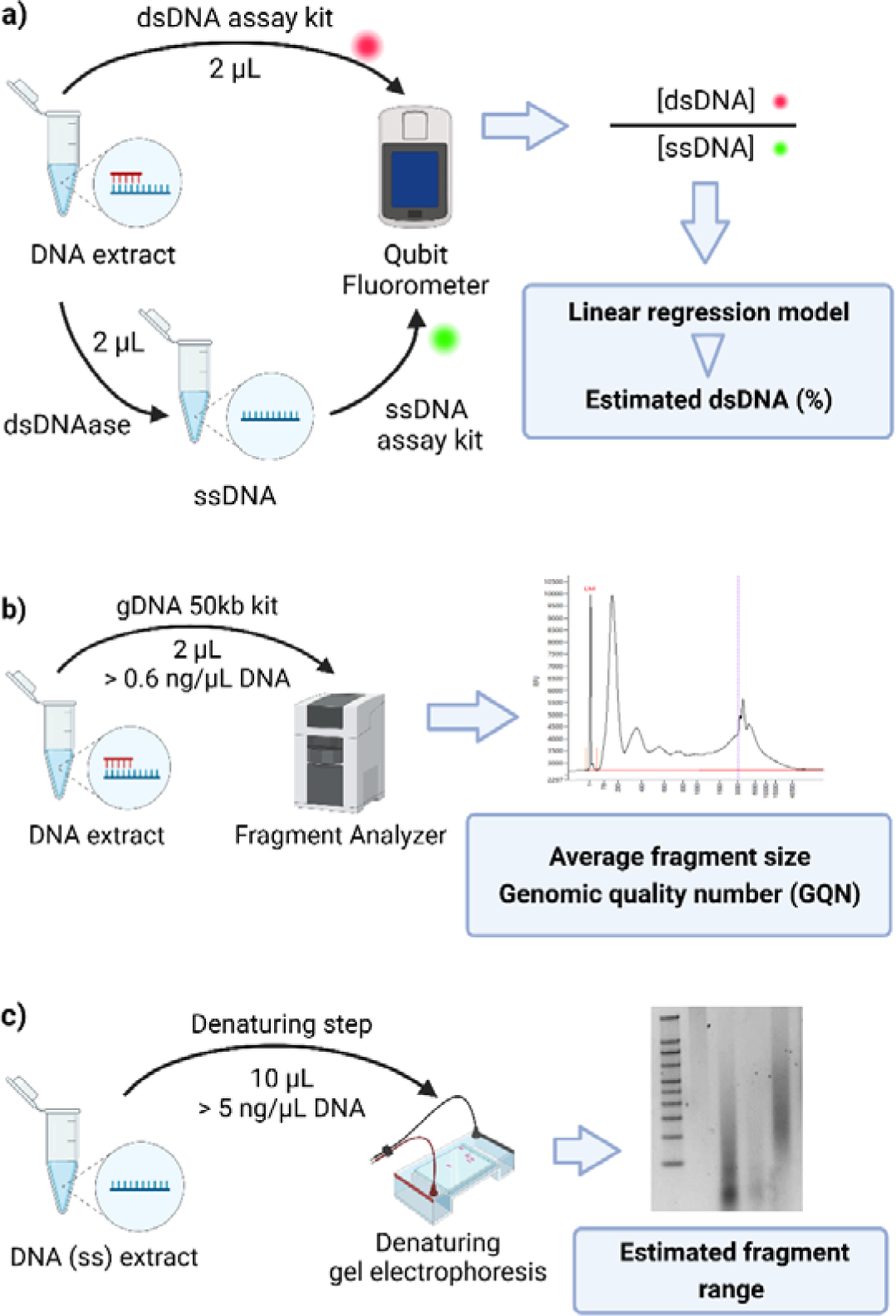
Overview of protocols and workflow for determining DNA quality. A) Generation of [dsDNA]/[ssDNA] for estimation of strandedness by a linear regression model. B) Analysis on Fragment Analyzer to generate the average DNA fragment size and genomic quality number (GQN) for samples with mainly dsDNA. c) Denaturing gel electrophoresis to estimate the fragment size range for samples with mainly ssDNA. The figure was created with BioRender.

#### 3.1.1 Development and validation of a protocol for determination of DNA strandedness

It is not trivial to identify a method for estimation of the proportion of dsDNA *versus* ssDNA in a sample, as most DNA quantification methods detect both dsDNA and ssDNA, although the signal intensities may differ [46, 47]. As we developed a fluorometric protocol based on the ratio of dsDNA and ssDNA Qubit measurements, we found that the Qubit dsDNA assay kit [38] is not exclusively providing signals from dsDNA but also from ssDNA, although in lower intensities. This was evident as the Qubit dsDNA assay kit generated fluorescent signals from pure ssDNA samples and from samples treated with dsDNases (S1 Table). Similarly, the Qubit ssDNA assay kit generates fluorescent signals from dsDNA in addition to ssDNA [40]. Thus, we included a dsDNase step prior to the ssDNA measurement. The ratios of the dsDNA and ssDNA values of reference samples with known proportions of dsDNA and ssDNA were plotted and a linear regression model was generated (y = 121.9x – 16.9, where x is the [dsDNA]/[ssDNA], R^2^ = 0.96) for prediction of DNA strandedness (Fig 2A). The model was validated against a second set of reference samples (n = 26), where all y values below 0 were set to 0% dsDNA and all y values above 100 were set to 100% dsDNA (Fig 2B). The mean absolute error (MAE) was 5.4 for all samples, but lower for samples with ≤ 25% dsDNA (MAE = 2.9) and higher for samples with ≥ 75% dsDNA (MAE = 7.6), indicative of a higher precision for samples with low proportions of dsDNA. The uncertainty of measurement suggests that when applying this regression model, percentages of dsDNA calculated to below 5 and above 95 should be presented as < 5% and >95%, respectively.

**Fig 2.**
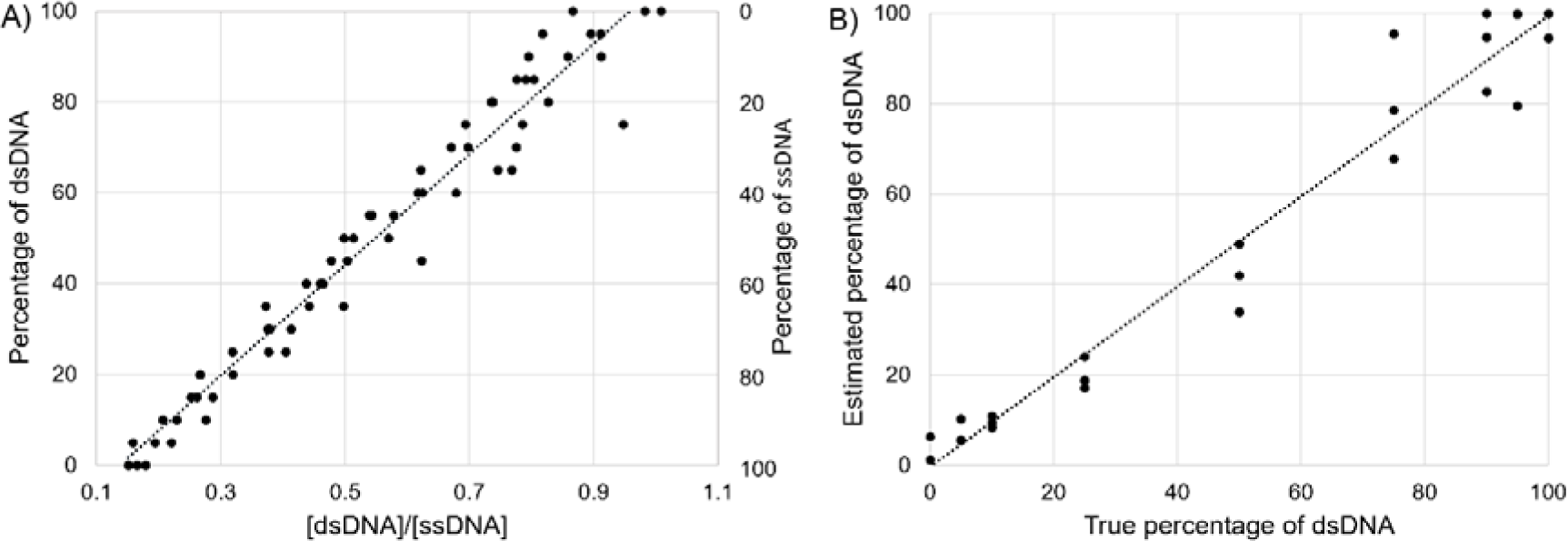
Estimation of DNA strandedness. A) A model to determine the percentage of dsDNA was generated using linear regression analysis, resulting in the linear equation y = 121.9x – 16.9, where x is [dsDNA]/[ssDNA] from Qubit measurements. The data is generated from reference samples with known proportions of dsDNA (left y axis) and ssDNA (right y axis), with total DNA concentrations of 0.5, 2 and 5 ng/µL (n = 63). B) Estimated versus true percentage of dsDNA, using the linear regression model (n = 26). All estimated values above 100% are set to 100%. R^2^ is 0.96.

The model was further tested on dilutions of saliva samples processed with two different DNA extraction protocols (QIAamp DNA Mini kit or Chelex protocol 2, n = 3 per protocol and dilution, S1 Table). [dsDNA]/[ssDNA] ratios were calculated for DNA concentrations 0.075 - 5 ng/µL where the mean predicted dsDNA percentage was not deemed significant between the dilutions (one-way ANOVA, p > 0.05). The LOQ of the method was determined as the lowest DNA concentration that generated quantifiable results with the Qubit dsDNA and ssDNA assay kits (2 µL of 0.075 ng/µL), *i.e*., providing a dsDNA/ssDNA concentration ratio. The precision of the method is indicated by the coefficients of variation (CV) calculated for the [dsDNA]/[ssDNA] of all replicates (17% for the QIAamp dilutions and 13% for the Chelex 2 dilutions). From this experiment, it was clear that saliva samples processed by QIAamp DNA Mini kit yielded higher amounts of dsDNA (above 95%) than samples processed by Chelex protocol 2 (13 – 30% dsDNA).

To estimate whether the majority of DNA in a sample is double- or single-stranded prior to library preparations, the sensitivity of the model we have developed is sufficient. The protocol is user-friendly, and the instrumentation affordable and convenient to implement. Another method to determine DNA strandedness has previously been presented by the National Institute of Standard and Technology (NIST). Their method relies on digital PCR and provides a resolution down to 2-3% ssDNA [48, 49].

#### 3.1.2 Evaluation of methods for determination of DNA fragmentation

First, we applied mock saliva samples processed by Chelex protocol 2 and QIAamp DNA Mini kit to compare the performance of Fragment Analyzer with Genomic DNA 50kb kit and TapeStation with High Sensitivity D5000 ScreenTape System. The LOQs were identical for the two systems, as dsDNA concentrations down to 0.6 ng/µL could be detected by both Fragment Analyzer and TapeStation. However, the TapeStation results for the Chelex processed extracts were considered unreliable as the TapeStation failed to detect the size markers for 50% of these samples. Further, the DNA concentrations given by TapeStation were not proportional to the known concentrations of the Chelex-processed samples (S1 Fig). Fragment Analyzer with Genomic DNA 50kb Kit was thus applied for further assessment of DNA fragment size.

For DNA dilutions (0.6 – 5 ng/µL, n = 3 per dilution) of saliva samples processed with QIAamp DNA Mini kit, no significant differences in average DNA fragment sizes were seen (24 ± 2.7 kb, one-way ANOVA: p > 0.05, S2 Table). The precision was 11% (CV), all replicates and dilutions taken together. The dilutions of Chelex-processed saliva samples showed an average DNA fragment size of 97 ± 13 bp, which most likely is an underestimation due to the high content of ssDNA in these samples. It is known that ssDNA migrates faster than dsDNA in a native gel such as the one used in the Fragment Analyzer, which will result in an incorrect fragment size determination of ssDNA [50, 51]. In addition to average fragment size, the genomic quality number (GQN) with a fragment size threshold of 10 kb was generated for each sample. GQN is a number from 0 – 10 that indicates the fraction of DNA with a larger size than the fragment size threshold, thus a GQN of 10 indicates that 100% of the DNA is above the fragment size threshold. For the DNA dilutions processed with either QIAmp DNA Mini Kit or Chelex protocol 2, there were no differences in the obtained GQN (one-way ANOVA: p > 0.05). However, the GQN differed significantly between samples processed with QIAamp DNA Mini kit and Chelex protocol 2 respectively (3.8 ± 0.1 and 0.08 ± 0.04, t-test: p < 0.001). Unprocessed human gDNA of high molecular weight analyzed with the Genomic DNA 50kb kit on Fragment Analyzer showed an average fragment size of 35 ± 7.4 kb and a GQN of 6.2 ± 0.31, serving as a reference to all processed samples.

It was concluded that analysis on Fragment Analyzer with Genomic DNA 50kb Kit provided reliable results regarding fragment size and GQN with dsDNA in concentrations between 0.6 – 5 ng/µL, but that results for samples with a higher percentage of ssDNA than dsDNA should be carefully interpreted. Preferably, alternative methods should be used for extracts known or expected to contain high proportions of ssDNA.

To assess the fragment size of ssDNA, we employed two different methods on DNA extracts processed with Chelex protocol 2, known from 3.1.1 to contain more than 70% ssDNA (S1 Table): TapeStation with High Sensitivity RNA ScreenTape System and denaturing agarose gel. Analysis on TapeStation with High Sensitivity RNA ScreenTape System gave slightly higher average fragment sizes (around 200 nt) compared to the Fragment Analyzer with Genomic DNA 50kb kit (around 100 bp, S1 Fig). However, single strands of DNA may form secondary sequence-dependent structures, and the migration rate is thus contingent on both size and conformation. Adding a denaturant to the gel, such as urea, results in single-stranded DNA without secondary structures and thus only the size of the DNA fragment will affect mobility. Analysis of undiluted DNA extracts processed by Chelex protocol 2 on denaturing gels resulted in a smear with denser regions over a broad size range (0.5 – 9 kb). It was difficult to estimate a precise fragment size from the smear, but it could be concluded that the DNA fragment sizes given by Fragment Analyzer (around 100 bp) and TapeStation with High Sensitivity RNA ScreenTape System (around 200 nt) were underestimations.

### 3.2 Impact on DNA quality of various DNA extraction protocols and sample types

The established methods (Fig 1) from section 3.1 were used to investigate how different DNA extraction methods and different mock casework samples (nasal secretion on swabs, blood, saliva on adhesive tapes, semen, and bone) affect the DNA quality in terms of strandedness and DNA fragmentation (Table 1). In addition, the DI values provided from qPCR analysis with PowerQuant System were generated for comparison to the DNA fragment sizes.

#### 3.2.1 DNA strandedness

The model for estimation of strandedness showed that the gDNA samples processed by different DNA extraction protocols provided very different percentages of dsDNA (Fig 3A, S3 Table). All non Chelex-based protocols generated more than 90% dsDNA, while all Chelex-based protocols gave less than 50% dsDNA. Chelex protocol 1 with a shorter heat incubation generated approximately 45% of dsDNA while Chelex protocol 2 and 3 and the differential lysis protocol with longer heat incubation steps gave less than 10% dsDNA. When the different mock casework samples were processed, we found that all samples extracted with QIAamp DNA Mini kit, Phenol protocol 1 and EZ1 Advanced XL contained more than 95% dsDNA (Fig 3A). Nasal secretion and saliva samples processed by Chelex protocol 1 – 3 showed similar percentages of dsDNA as the gDNA (around 40% dsDNA with protocol 1 and less than 15% dsDNA with protocol 2 and 3). Semen samples extracted with the Chelex-based differential lysis protocol also gave a low percentage of dsDNA (< 20%). DNA extracted from blood remained mainly double stranded for the Chelex protocols 1 and 2 but not for Chelex protocol 3 which resulted in around 20% dsDNA.

**Fig 3.**
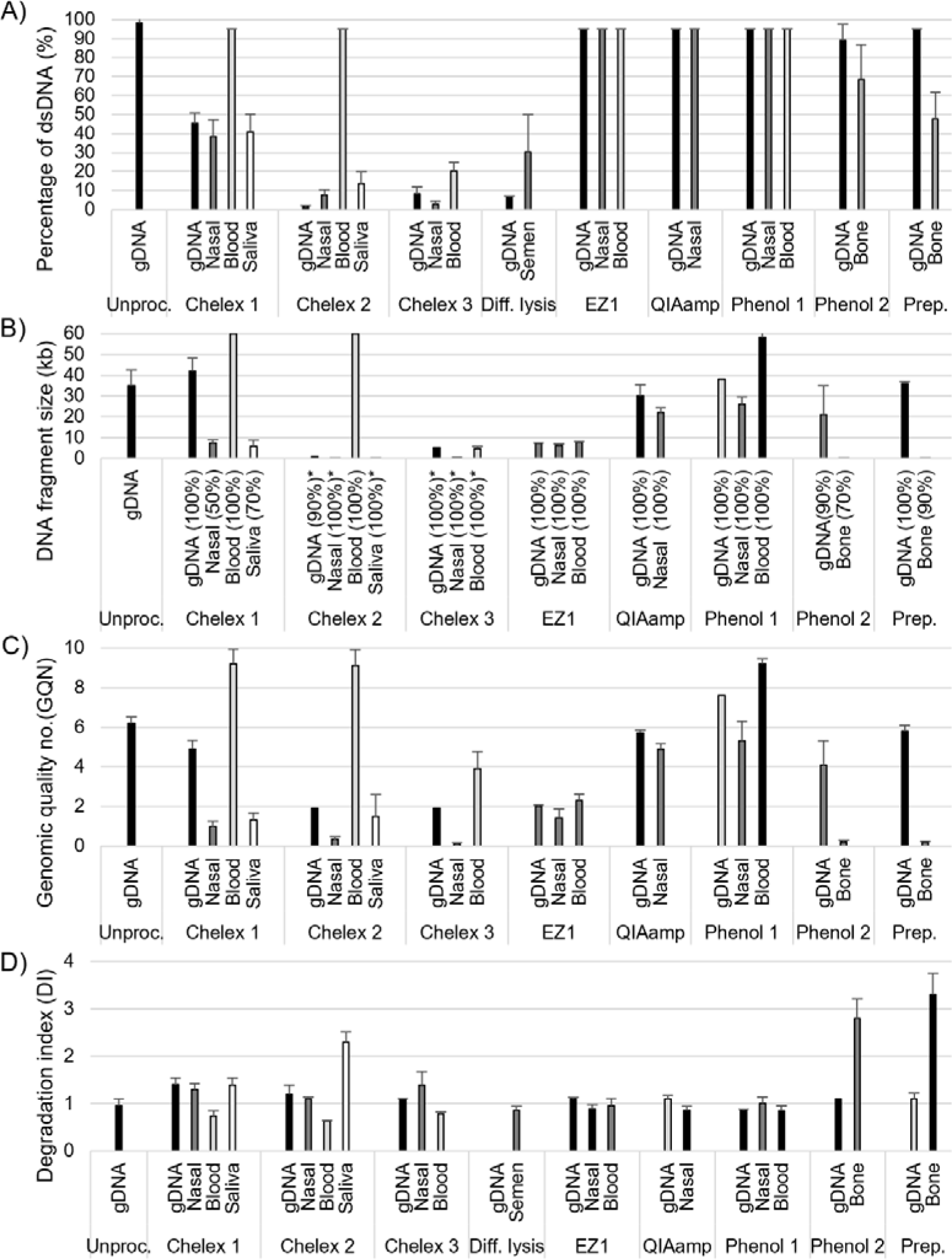
The effects of different DNA extraction protocols and sample types on DNA quality. The effects on A) DNA strandedness, B) average DNA fragment size, C) genomic quality number (GQN), and D) degradation index (DI) as given by PowerQuant System qPCR are shown as average values ± standard deviations (n = 2-9, see S2 Table for exact number). DNA strandedness is presented as the approximate proportion of dsDNA (%). DNA fragmentation is presented as the average fragment size of the major DNA peak as given by the Fragment Analyzer. The percentage of total DNA in the major peak is presented in parenthesis. Asterisk (*) indicates that the fragment sizes may be underestimated as a cause of higher gel migration speed of single-stranded DNA. The effect of the extraction protocol alone is given by the strandedness/fragment size of gDNA. Additional effects of sample types are given for nasal secretion on cotton swab, blood, saliva on adhesive tapes, semen, and bone. The fragment size and GQN are not given for samples processed with differential lysis due to failed detection. Unproc: Unprocessed samples, Diff. lysis: Differential lysis, Prep: PrepFiler Express.

Others have previously shown that Chelex-based extraction protocols generate single-stranded DNA and that the proportion of ssDNA increases with longer heat incubation [52-55]. In addition to heat exposure at 100°C, the alkalinity of the Chelex suspension (pH 10-11) causes denaturing of DNA during the Chelex procedure [46, 52]. Additionally, Chelex chelates magnesium ions which help to counteract the electrostatic repulsion of the DNA strands, further promoting ssDNA. However, subsequent DNA purification of Chelex extracts including exchange of storage buffer to phosphate buffered saline (PBS) with pH 8, was not sufficient for renaturation of the DNA (S2_File_A).

Bone extracted with PrepFiler or Phenol protocol 2 resulted in around 50-70% dsDNA, which may be attributed to initial poor DNA quality of the bone samples [3, 56]. This is supported by comparing the results to the gDNA processed with PrepFiler and Phenol protocol 2, which remained double stranded (> 90% dsDNA). The Phenol protocol 2 gave a slightly higher percentage of dsDNA compared to PrepFiler Express. This is concordant with previous studies describing that the choice of extraction method influences DNA quality from bone samples [4, 57, 58].

Re-analysis of all mock casework samples after one year showed that the percentage of dsDNA remained rather stable (S4 Table). None of the sample groups showed an increased proportion of ssDNA after one year in -20 °C.

#### 3.2.2 DNA fragmentation

DNA fragmentation was assessed by fragment size and peak width of the two major peaks in the Fragment Analyzer electropherogram, along with the GQN with a threshold of 10 kb (Figs 3B-C and 4, S3 Table). Undiluted extracts containing a majority of ssDNA were analyzed on denaturing gels to assess the fragment sizes of ssDNA (S2 Fig).

Analysis of gDNA samples processed by the different DNA extraction protocols revealed that Chelex protocol 1, QIAamp DNA Mini kit, Phenol protocol 1 and PrepFiler Express resulted in large DNA fragments (30 – 42 kb), similar to the unprocessed gDNA (35 ± 7.4 kb, Fig 3B). Phenol protocol 2 gave fragment sizes around 21 kb and EZ1 Advanced XL yielded DNA fragments around 7 kb. gDNA processed with Chelex protocol 2 and 3 provided DNA fragments of approximately 1 and 5 kb, respectively, when analyzed on Fragment Analyzer. No DNA peaks were detected in gDNA samples processed with differential lysis, possibly due to DNA amounts below the detection limit.

Nasal secretion samples processed with QIAamp DNA Mini kit and Phenol protocol 1 yielded slightly smaller DNA fragments compared to the gDNA, with average fragment sizes of 22 ± 2.1 kb and 26 ± 3.8 kb, respectively. Nasal secretion and blood samples processed with the EZ1 Advanced XL harbored DNA fragments of around 7 kb, similar to the gDNA samples. Such limited range of DNA fragment sizes may depend on the concentration and size of magnetic beads in the EZ1 method, factors known to dictate the DNA size binding preference [59]. Chelex protocol 1 gave one peak with average DNA fragments of 7.5 ± 1.7 kb (nasal secretion) and 5.6 ± 3.4 kb (saliva), and one additional peak with fragments around 200 bp. The results from denaturing gel electrophoresis for these samples showed a constant smear ranging from 1-2 to above 9 kb. In comparison to the gDNA processed with Chelex protocol 1, the DNA from nasal secretion on swabs and saliva on adhesive tapes processed with the same method appears to be much more fragmented. The majority of DNA from nasal secretion and saliva samples processed with Chelex protocol 2 and 3 were shown to be highly fragmented (< 300 bp) by the Fragment Analyzer, but the denaturing gels showed a smear with a denser region between 0.5 – 3 kb.

When bone samples were processed by PrepFiler Express and Phenol protocol 2, the DNA was shown to be highly fragmented (around 200 bp with Fragment Analyzer and < 1 kb with denaturing gels). This is not surprising, as bone casework samples have been exposed to the decay process beginning at the moment of death, where released enzymes and microbial activity rapidly cause degradation of DNA [3, 56]. The analysis of semen extracted with a differential lysis method yielded fluorescence spikes in the Fragment Analyzer and these results could not be used. Semen samples on denaturing gels showed a smear with a denser region and a thin band > 9 kb (S2 Fig).

DNA derived from blood consisted of large DNA fragments regardless of the protocol applied. All DNA from the blood samples processed with Chelex protocol 1 and 2 had an average fragment size of above 60 kb, while blood samples processed with Chelex 3 and Phenol 1 gave average fragment sizes of 4.8 ± 1.4 kb and 58 ± 3.1 kb, respectively. This is remarkable as all other biological sample types gave smaller single-stranded DNA fragments when processed with the Chelex protocols. We speculated that the addition of EDTA, added as an anti-coagulant to the blood in the mock casework samples, counteracted fragmentation and denaturation of DNA due to its capacity to chelate metal cations required as cofactors for DNases. It has been shown that addition of EDTA to blood samples reduces DNA degradation due to its inhibition of DNase activity [60, 61]. As we examined the DNA strandedness and fragmentation of blood samples extracted with Chelex protocol 1 with and without the addition of EDTA (n = 3), the results showed that all blood samples gave over 95% dsDNA of high fragment size (> 60 kb), regardless of EDTA addition. This indicates that EDTA is not responsible for counteracting fragmentation and denaturation of DNA during the extraction process of blood samples.

The GQN indicates the fraction of DNA of larger size than the fragment size threshold and may be used as a complement to the average DNA fragment size (Figs 3C and 5A). GQN and DNA fragment size are derived from the same Fragment Analyzer data but while the GQN provides one value from 1 to 10 for the total DNA amount, the average fragment size is given for each DNA peak in the electropherogram (Fig 4). If the GQN with a fragment size threshold of 10 kb is approaching 10, the vast majority of DNA consists of fragments larger than 10 kb and is considered “genomic” in the DNA shearing protocols of Covaris [62], an ultrasonicator instrument commonly used to shear DNA prior to library preparations. For samples with smaller DNA fragments than 10 kb, Covaris recommends setting up a time dose response experiment for determining appropriate shearing times.

**Fig 4.**
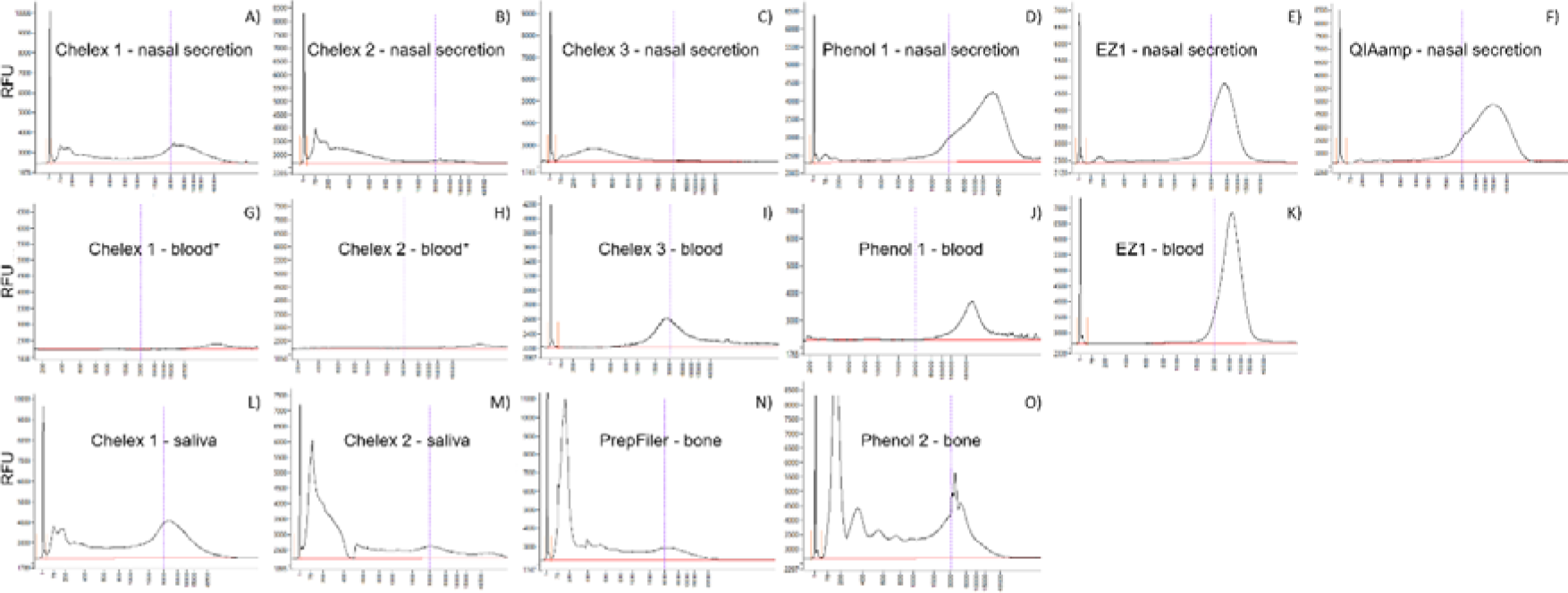
Electropherograms generated by the Fragment Analyzer. Representative DNA fragment sizes for nasal secretion samples (A-F), blood samples (G-K), and saliva and bone samples (L-O) extracted using Chelex protocols 1-3, Phenol protocol 1-2, BioRobot EZ1 Advanced XL, QIAamp DNA Mini kit, and PrepFiler Express BTA Forensic DNA extraction kit. Electropherograms are shown for 5 ng/µL DNA extracts, except for blood samples extracted with Chelex protocol 1-2 (*) which are shown for 2 ng/µL dilutions. The x axes show DNA fragment size (bp) in logarithmic scale (log 10). The y axes show relative fluorescence units (RFU). It should be noted that the scales on the y axes differ between the graphs. The vertical lines in the graphs indicate the position of DNA fragments of 3 kb.

**Fig 5.**
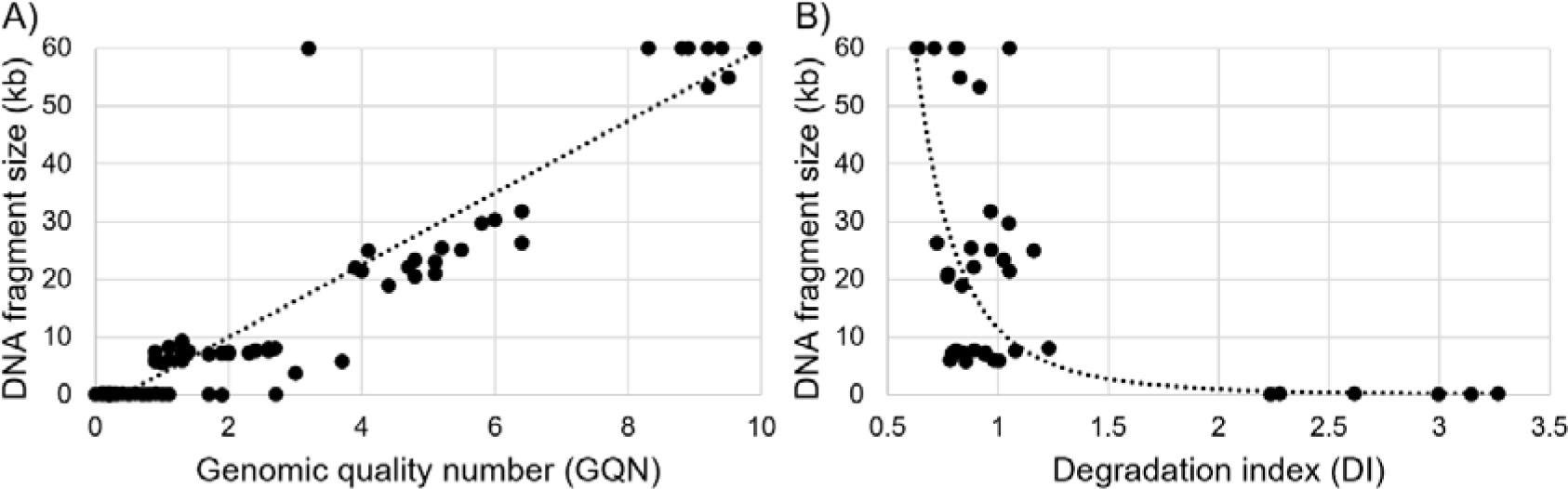
DNA fragment size plots. The average DNA fragment sizes are plotted against A) the genomic quality numbers (GQN) with a threshold of 10 kb, and B) the degradation indices (DI) provided by PowerQuant qPCR. Data on y-axis shows the average DNA fragment size of the major peak given by Fragment Analyzer. Data on GQN includes all mock casework samples (n = 95) and data on DI includes all mock casework samples with more than 60% double stranded DNA (n = 38).

A common method for assessing DNA degradation in forensics is to compare the amplification of one small and one large DNA target in qPCR [63, 64]. With the PowerQuant System that was used in our study, a DI above 2 signifies DNA degradation of a degree that may be critical to forensic DNA profiling [63, 65]. In our study, the relationship between the DI and average fragment size of the major peak for all samples harboring more than 60% dsDNA is best described by a non-linear power regression model (R^2^ = 0.77, p < 0.0001, n = 38, Fig 5B). According to this model, the average DNA fragment size must be around 1 kb or smaller to generate a DI of 2 or above. Thus, applying the DI to predict average DNA fragment sizes larger than 1 kb is not very useful, as the DI values are approximately 1 regardless of a DNA fragment size of 5 or 60 kb (Figs 3B, 3D and 5). In accordance with our data, it has previously been shown that the relationship between DI values and DNA fragment sizes is non-linear [66] and that the DI values indeed increase as samples are exposed to DNA degrading treatments such as sonication and UV-C radiation [65, 67]. The DI is undoubtably a valuable tool to predict DNA degradation prior to forensic DNA profiling that relies on amplification of target sequences smaller than 500 bp, although substantial deviations between the DI value and the impact DNA degradation has on the forensic DNA profiles also have been reported [68, 69]. For estimating DNA fragment sizes larger than 1 kb, as required for correct shearing parameters prior to library preparation, assessing the DI is not an optimal method.

Re-analysis of all samples after one year on Fragment Analyzer showed that the average DNA fragment sizes were relatively stable, although small but statistically significant reductions in fragment size were seen for nasal secretion samples processed with Chelex protocol 2 and for bone processed with Phenol protocol 2 (S4 Table). An increase in measured fragment size was seen for blood processed with Chelex protocol 3 (from 4.8 ± 1.0 to >60 kb, p < 0.01), which is difficult to explain. We speculated that the presence of blood may disturb the fluorescent signal from the DNA-binding dye applied in the analysis methods, causing this anomaly. For example, it has been shown that immunoglobulin G binds with a high affinity to ssDNA and that hemoglobin and hematin can quench the fluorescent signal in qPCR [70]. We thus performed an experiment where whole blood was added to samples with dsDNA and ssDNA extracts of known concentrations, but we found no effect of blood on the Qubit signal (S1_File_B).

### 3.3 Determination of DNA quality of forensic casework samples

45 forensic casework samples of various types were processed with different DNA extraction protocols and assessed for DNA strandedness and fragmentation (Table 2).

**Table 2.**
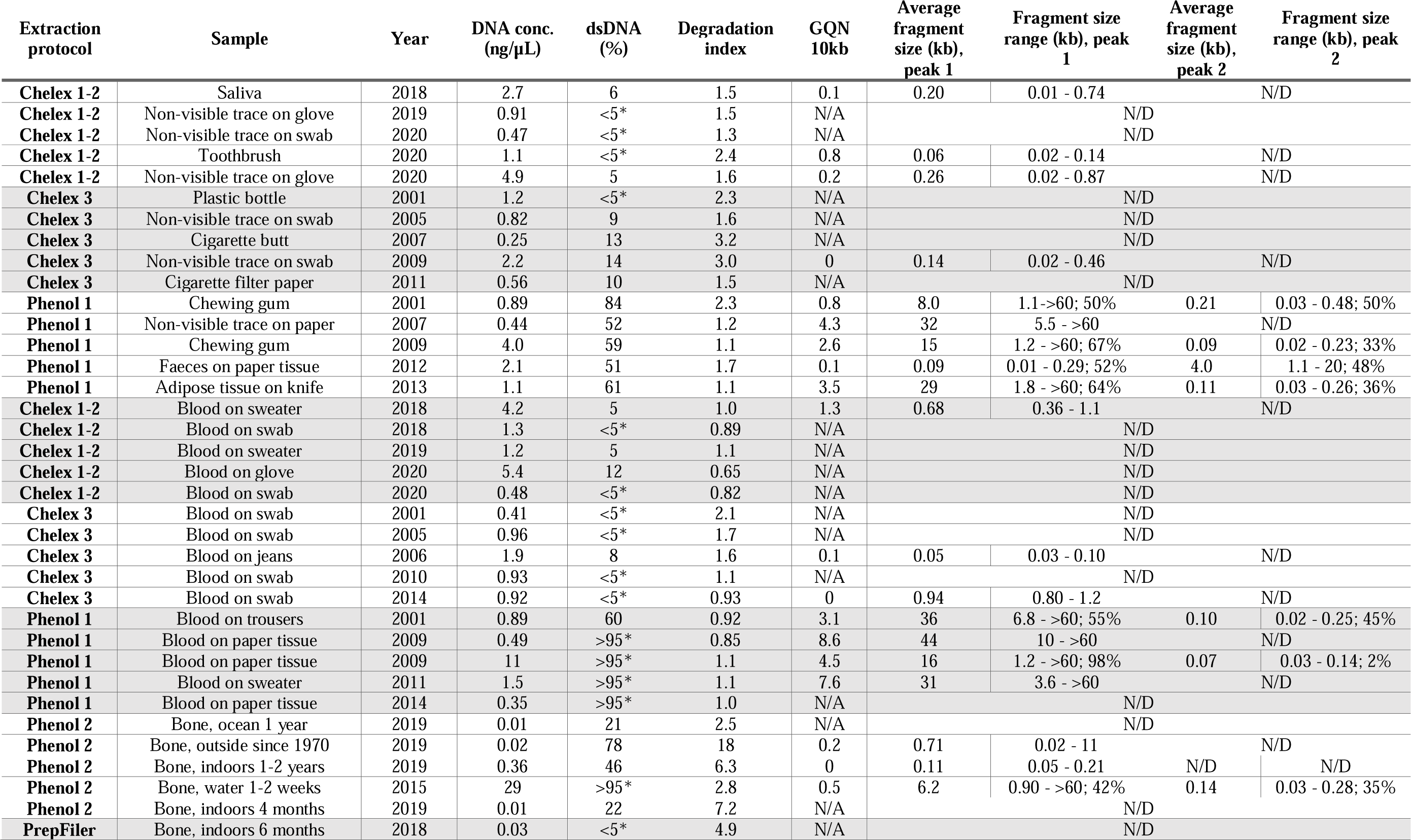
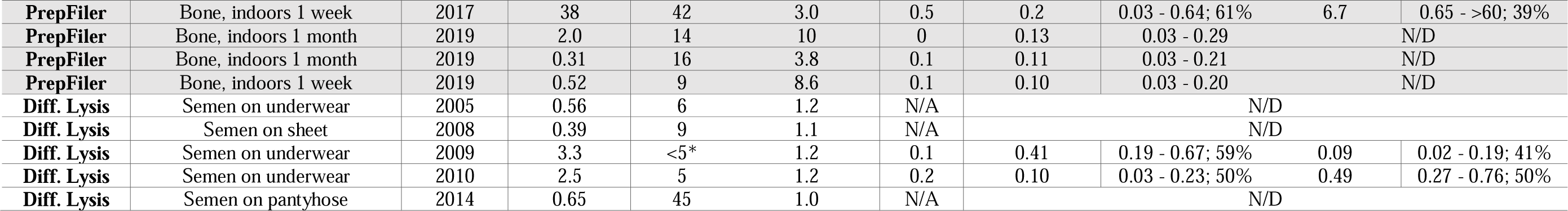
DNA quality of forensic casework samples. The table includes information on DNA extraction protocols, sample description and year of collection. The following data are presented: DNA concentration (ng/µL) obtained from qPCR with PowerQuant, percentage of dsDNA, degradation index (given by qPCR, PowerQuant), the GQN_10kb_ and average fragment size and width for peak 1 and 2 (kb) including the percentages of total DNA the peaks constitute (given by Fragment Analyzer). N/D: not detected, N/A: not available. Asterisk (*) indicates that the calculated percentage of dsDNA were below 5 or above 95, and thus approximated to < 5% or >95%.

| Extraction protocol | Sample | Year | DNA conc. (ng/μL) | dsDNA (%) | Degradation index | GQN 10kb | Average fragment size (kb), peak 1 | Fragment size range (kb), peak 1 | Average fragment size (kb), peak 2 | Fragment size range (kb), peak 2 |
| --- | --- | --- | --- | --- | --- | --- | --- | --- | --- | --- |
| **Chelex 1-2** | Saliva | 2018 | 2.7 | 6 | 1.5 | 0.1 | 0.20 | 0.01 - 0.74 |  | N/D |
| **Chelex 1-2** | Non-visible trace on glove | 2019 | 0.91 | <5* | 1.5 | N/A |  | N/D |  |  |
| **Chelex 1-2** | Non-visible trace on swab | 2020 | 0.47 | <5* | 1.3 | N/A |  | N/D |  |  |
| **Chelex 1-2** | Toothbrush | 2020 | 1.1 | <5* | 2.4 | 0.8 | 0.06 | 0.02 - 0.14 |  | N/D |
| **Chelex 1-2** | Non-visible trace on glove | 2020 | 4.9 | 5 | 1.6 | 0.2 | 0.26 | 0.02 - 0.87 |  | N/D |
| **Chelex 3** | Plastic bottle | 2001 | 1.2 | <5* | 2.3 | N/A |  | N/D |  |  |
| **Chelex 3** | Non-visible trace on swab | 2005 | 0.82 | 9 | 1.6 | N/A |  | N/D |  |  |
| **Chelex 3** | Cigarette butt | 2007 | 0.25 | 13 | 3.2 | N/A |  | N/D |  |  |
| **Chelex 3** | Non-visible trace on swab | 2009 | 2.2 | 14 | 3.0 | 0 | 0.14 | 0.02 - 0.46 |  | N/D |
| **Chelex 3** | Cigarette filter paper | 2011 | 0.56 | 10 | 1.5 | N/A |  | N/D |  |  |
| **Phenol 1** | Chewing gum | 2001 | 0.89 | 84 | 2.3 | 0.8 | 8.0 | 1.1->60; 50% | 0.21 | 0.03 - 0.48; 50% |
| **Phenol 1** | Non-visible trace on paper | 2007 | 0.44 | 52 | 1.2 | 4.3 | 32 | 5.5 - >60 |  | N/D |
| **Phenol 1** | Chewing gum | 2009 | 4.0 | 59 | 1.1 | 2.6 | 15 | 1.2 - >60; 67% | 0.09 | 0.02 - 0.23; 33% |
| **Phenol 1** | Faeces on paper tissue | 2012 | 2.1 | 51 | 1.7 | 0.1 | 0.09 | 0.01 - 0.29; 52% | 4.0 | 1.1 - 20; 48% |
| **Phenol 1** | Adipose tissue on knife | 2013 | 1.1 | 61 | 1.1 | 3.5 | 29 | 1.8 - >60; 64% | 0.11 | 0.03 - 0.26; 36% |
| **Chelex 1-2** | Blood on sweater | 2018 | 4.2 | 5 | 1.0 | 1.3 | 0.68 | 0.36 - 1.1 |  | N/D |
| **Chelex 1-2** | Blood on swab | 2018 | 1.3 | <5* | 0.89 | N/A |  | N/D |  |  |
| **Chelex 1-2** | Blood on sweater | 2019 | 1.2 | 5 | 1.1 | N/A |  | N/D |  |  |
| **Chelex 1-2** | Blood on glove | 2020 | 5.4 | 12 | 0.65 | N/A |  | N/D |  |  |
| **Chelex 1-2** | Blood on swab | 2020 | 0.48 | <5* | 0.82 | N/A |  | N/D |  |  |
| **Chelex 3** | Blood on swab | 2001 | 0.41 | <5* | 2.1 | N/A |  | N/D |  |  |
| **Chelex 3** | Blood on swab | 2005 | 0.96 | <5* | 1.7 | N/A |  | N/D |  |  |
| **Chelex 3** | Blood on jeans | 2006 | 1.9 | 8 | 1.6 | 0.1 | 0.05 | 0.03 - 0.10 |  | N/D |
| **Chelex 3** | Blood on swab | 2010 | 0.93 | <5* | 1.1 | N/A |  | N/D |  |  |
| **Chelex 3** | Blood on swab | 2014 | 0.92 | <5* | 0.93 | 0 | 0.94 | 0.80 - 1.2 |  | N/D |
| **Phenol 1** | Blood on trousers | 2001 | 0.89 | 60 | 0.92 | 3.1 | 36 | 6.8 - >60; 55% | 0.10 | 0.02 - 0.25; 45% |
| **Phenol 1** | Blood on paper tissue | 2009 | 0.49 | >95* | 0.85 | 8.6 | 44 | 10 - >60 |  | N/D |
| **Phenol 1** | Blood on paper tissue | 2009 | 11 | >95* | 1.1 | 4.5 | 16 | 1.2 - >60; 98% | 0.07 | 0.03 - 0.14; 2% |
| **Phenol 1** | Blood on sweater | 2011 | 1.5 | >95* | 1.1 | 7.6 | 31 | 3.6 - >60 |  | N/D |
| **Phenol 1** | Blood on paper tissue | 2014 | 0.35 | >95* | 1.0 | N/A |  | N/D |  |  |
| **Phenol 2** | Bone, ocean 1 year | 2019 | 0.01 | 21 | 2.5 | N/A |  | N/D |  |  |
| **Phenol 2** | Bone, outside since 1970 | 2019 | 0.02 | 78 | 18 | 0.2 | 0.71 | 0.02 - 11 |  | N/D |
| **Phenol 2** | Bone, indoors 1-2 years | 2019 | 0.36 | 46 | 6.3 | 0 | 0.11 | 0.05 - 0.21 | N/D | N/D |
| **Phenol 2** | Bone, water 1-2 weeks | 2015 | 29 | >95* | 2.8 | 0.5 | 6.2 | 0.90 - >60; 42% | 0.14 | 0.03 - 0.28; 35% |
| **Phenol 2** | Bone, indoors 4 months | 2019 | 0.01 | 22 | 7.2 | N/A |  | N/D |  |  |
| **PrepFiler** | Bone, indoors 6 months | 2018 | 0.03 | <5* | 4.9 | N/A |  | N/D |  |  |
| **PrepFiler** | Bone, indoors 1 week | 2017 | 38 | 42 | 3.0 | 0.5 | 0.2 | 0.03 - 0.64; 61% | 6.7 | 0.65 - >60; 39% |
| **PrepFiler** | Bone, indoors 1 month | 2019 | 2.0 | 14 | 10 | 0 | 0.13 | 0.03 - 0.29 |  | N/D |
| **PrepFiler** | Bone, indoors 1 month | 2019 | 0.31 | 16 | 3.8 | 0.1 | 0.11 | 0.03 - 0.21 |  | N/D |
| **PrepFiler** | Bone, indoors 1 week | 2019 | 0.52 | 9 | 8.6 | 0.1 | 0.10 | 0.03 - 0.20 |  | N/D |
| **Diff. Lysis** | Semen on underwear | 2005 | 0.56 | 6 | 1.2 | N/A |  |  |  | N/D |
| **Diff. Lysis** | Semen on sheet | 2008 | 0.39 | 9 | 1.1 | N/A |  |  |  | N/D |
| **Diff. Lysis** | Semen on underwear | 2009 | 3.3 | <5* | 1.2 | 0.1 | 0.41 | 0.19 - 0.67; 59% | 0.09 | 0.02 - 0.19; 41% |
| **Diff. Lysis** | Semen on underwear | 2010 | 2.5 | 5 | 1.2 | 0.2 | 0.10 | 0.03 - 0.23; 50% | 0.49 | 0.27 - 0.76; 50% |
| **Diff. Lysis** | Semen on pantyhose | 2014 | 0.65 | 45 | 1.0 | N/A |  |  |  | N/D |

All samples processed with Chelex-based protocols exhibited dsDNA percentages below 14% except for one semen sample that contained 45% dsDNA. Extraction with Phenol protocol 1 gave a dsDNA content ranging from 51 to 84% for nasal secretion samples and from 60 to above 95% for blood samples. The percentage of dsDNA in bone samples ranged from 21 to above 95% when processed with Phenol protocol 2 and from 2 to 42% when PrepFiler Express was applied. The proportions of dsDNA in the forensic casework extracts were lower compared to the mock casework samples compared on group level (S5 Table). This implies that apart from the sample type (nasal secretion on swabs, blood, saliva on adhesive tapes, semen, and bone) and the applied DNA extraction protocol, additional factors such as storage time or environmental conditions at the sampling site may affect the strandedness. The most notable difference was seen for blood samples extracted with Chelex protocol 1-2, with mock casework samples containing >95% dsDNA and forensic casework samples around 5% dsDNA. Considering that the re-analysis of the mock casework extracts after one year in -20°C resulted in similar or higher percentages of dsDNA in the Chelex-processed samples (S4 Table), it is not plausible that DNA in the forensic casework extracts goes from double-stranded to almost completely single-stranded after a few years in storage. Rather, we hypothesized that the high percentage of dsDNA in blood-derived mock casework samples could be due to higher amounts of processed blood or the immediate freezing of the mock casework samples after collection. Forensic casework samples may originate from less than a few microliters blood and the time between deposition and sample collection may range from days to years. We assessed these differences experimentally and found that regardless of the amount of blood in the Chelex-processed samples, the DNA remained double stranded (S1_File_C). When processing blood stains collected at different time points after deposition (from 0 days to 8 weeks), we found no substantial differences in DNA strandedness between any of the time points (S1_File_D). It has previously been reported that DNA from blood samples stored at room temperature shows a higher level of degradation compared to directly processed samples [68, 71, 72], and it is common knowledge that high molecular weight DNA best can be isolated from fresh tissue. However, the molecular mechanisms explaining the differences in DNA qualities we observed in forensic and mock casework samples respectively, remains to be elucidated.

The Fragment Analyzer was able to detect DNA peaks in 9 of 25 samples extracted with the Chelex-based protocols. These samples all exhibited small DNA fragments (< 1 kb, Table 2), but as previously stated the fragment size of ssDNA is likely underestimated due to a faster gel migration speed compared to dsDNA. Unfortunately, the low volumes of available forensic casework samples did not allow for analysis on denaturing gels. Forensic casework samples derived from non-visible traces, saliva or faeces that were processed with Phenol protocol 1 harbored DNA fragments ranging 0.09 - 32 kb while blood-derived samples revealed larger fragment sizes ranging 16 - 44 kb. In accordance, DNA extracted from non-visible DNA traces, saliva and faeces revealed higher DI values (1.82 ± 0.64) compared to blood samples (1.12 ± 0.37, p = 0.002, n = 15). Most forensic casework samples derived from bone had a high degree of fragmented DNA (< 1kb), in line with the PowerQuant DI values (2.5 – 18).

A comparison of the fragment sizes and DI values between mock and forensic casework samples suggests a higher level of DNA degradation in forensic casework samples for most sample groups (S5 Table). This suggests that DNA fragmentation, similar to DNA strandedness, is also affected by factors such as storage and original sample conditions, in addition to sample type and applied DNA extraction protocol. While numerous studies have shown that Chelex-based extraction protocols are cost-effective with similar or better performance compared to commercially available kits [6, 52, 73-78], concerns have been raised whether Chelex-based extracts provide adequate DNA stability for long-term storage [79]. The high pH (10-11) of the Chelex solution has been shown to decrease over time [80], and the chelating effect of the Chelex resin, removing metal cations required for DNase activity, may also decline over time. However, we find no clear indications that DNA fragmentation increases in Chelex-processed samples during storage for 1 year (S4 Table). In agreement, other studies have shown that storage up to 3 years and several freeze-thaw cycles of DNA samples generated from Chelex-based extractions showed no negative effects on DNA concentrations or forensic STR profile quality [78, 81]. In the past, storage conditions of forensic samples have been adjusted and validated primarily to preserve the DNA for an eventual re-analysis of DNA profiles based on STRs [81]. Although PCR-based generation of STR profiles is theoretically indifferent to whether the DNA is single- or double-stranded from the start [54], slightly fragmented single-stranded DNA molecules may provide an advantage in PCR due to easier accessibility for the primers to bind to its template DNA. In whole genome DNA library preparations however, it is crucial to know whether the DNA is single- or double-stranded. For example, the ThruePLEX kit, designed for dsDNA, was employed in two Swedish case studies [23, 24], wherein the later Chelex-processed samples were assessed and repeated analyses were needed to obtain useful results. For samples with a majority of ssDNA, there are more suitable kits available, for instance the SRSLY kit from Claret Bioscience [82]. The insight on typical DNA characteristics of different sample types provides important knowledge that may be used for guidance to which library preparation kit to apply. By also knowing the initial level of DNA fragmentation in a sample, excessive DNA shearing may be avoided through proper adjustments of shearing parameters prior to library preparation. The future prospective of applying SNP and WGS, or targeted hybridization capture sequencing, on degraded DNA of low quantity is undeniably a potent tool.

## 4. Conclusions

A fluorometry-based model for estimation of DNA strandedness was developed and validated to give reliable results for low amounts of DNA (0.15 ng). Analysis of DNA fragment sizes using Fragment Analyzer was evaluated and considered useful for samples with mostly double stranded DNA and a total DNA concentration ≥0.6 ng/µL. However, for samples with a high content of single-stranded DNA, the fragment size was underestimated and must be interpreted with caution. DNA extracts from forensic mock and forensic casework samples were assessed for DNA strandedness and fragmentation. The results show that the choice of DNA extraction protocol highly influences whether the DNA comes out as single- or double-stranded, as well as the level of DNA fragmentation. It was also found that DNA quality in terms of DNA strandedness and fragmentation was dependent on the sample type. DNA derived from bone samples was often at least partly single-stranded and fragmented in contrast to other forensic samples extracted with a phenol-based protocol. Interestingly, DNA derived from blood in mock casework samples remained double stranded and of high molecular weight compared to DNA originating from saliva and nasal secretion, when the same DNA extraction protocols were applied. However, forensic casework samples processed by Chelex-based protocols showed single-stranded and highly fragmented DNA. The results of this study have provided knowledge of how different DNA extraction methods influence DNA quality in typical forensic DNA samples, which may be useful for future selections of appropriate DNA extraction protocols. The established workflow to estimate DNA quality in “old” samples such as from cold cases and unidentified human remains will be valuable for prospective sequencing-based high-density SNP genotyping, as it enables the choice of an appropriate library preparation kit with DNA shearing parameters adjusted to the specific fragmentation level. Such workflows and knowledge are also applicable to other fields, such as clinical genetics and archaeology.

## 5. Acknowledgement

We would like to express our gratitude to the donors of cell material that made this study possible.

## Notes

### Competing Interest Statement

The authors have declared no competing interest.

