## Supplementary figures and images for "Assessment of DNA quality for whole genome library preparation"

### S1_Fig.tif

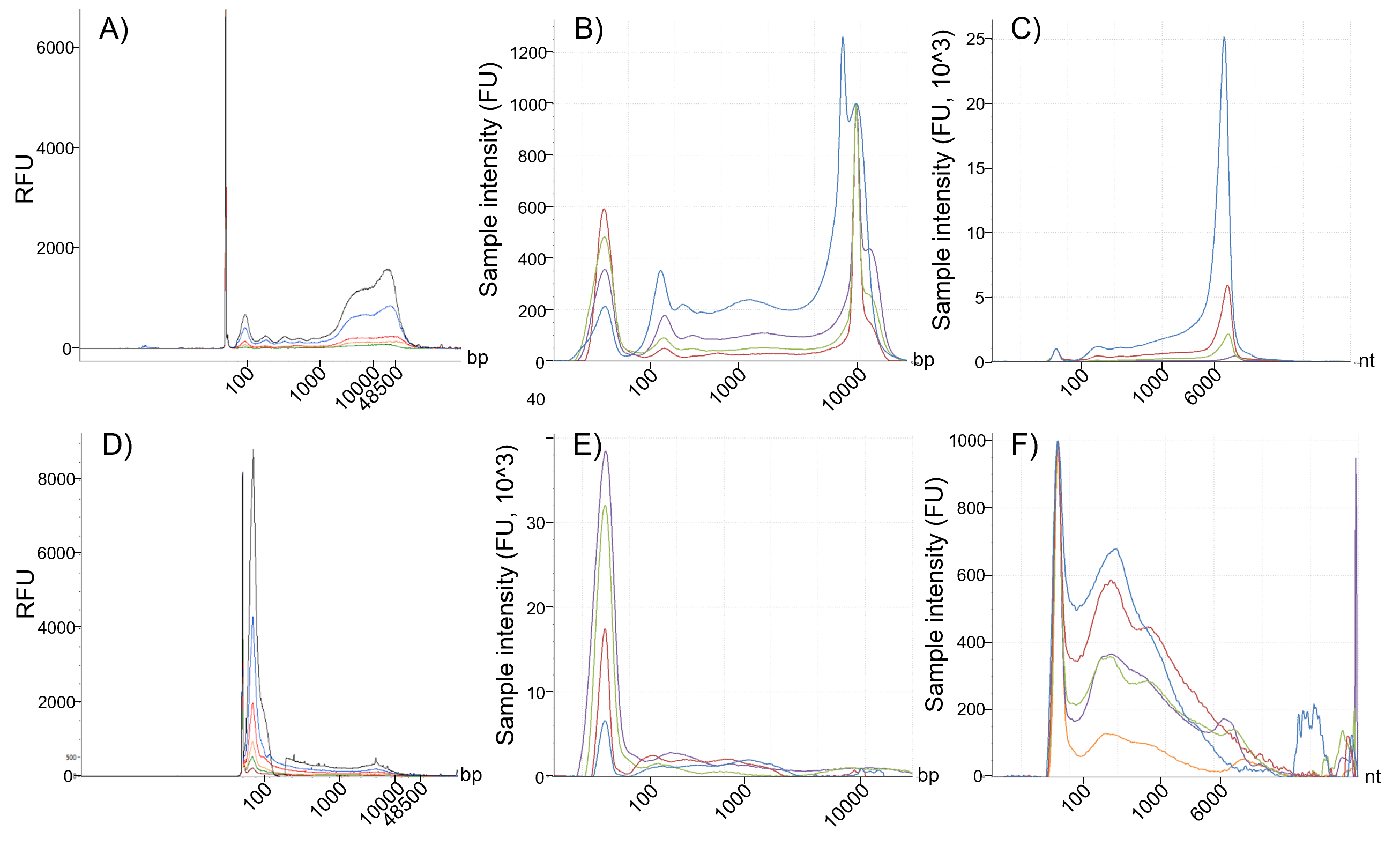

### S2_Fig.tif

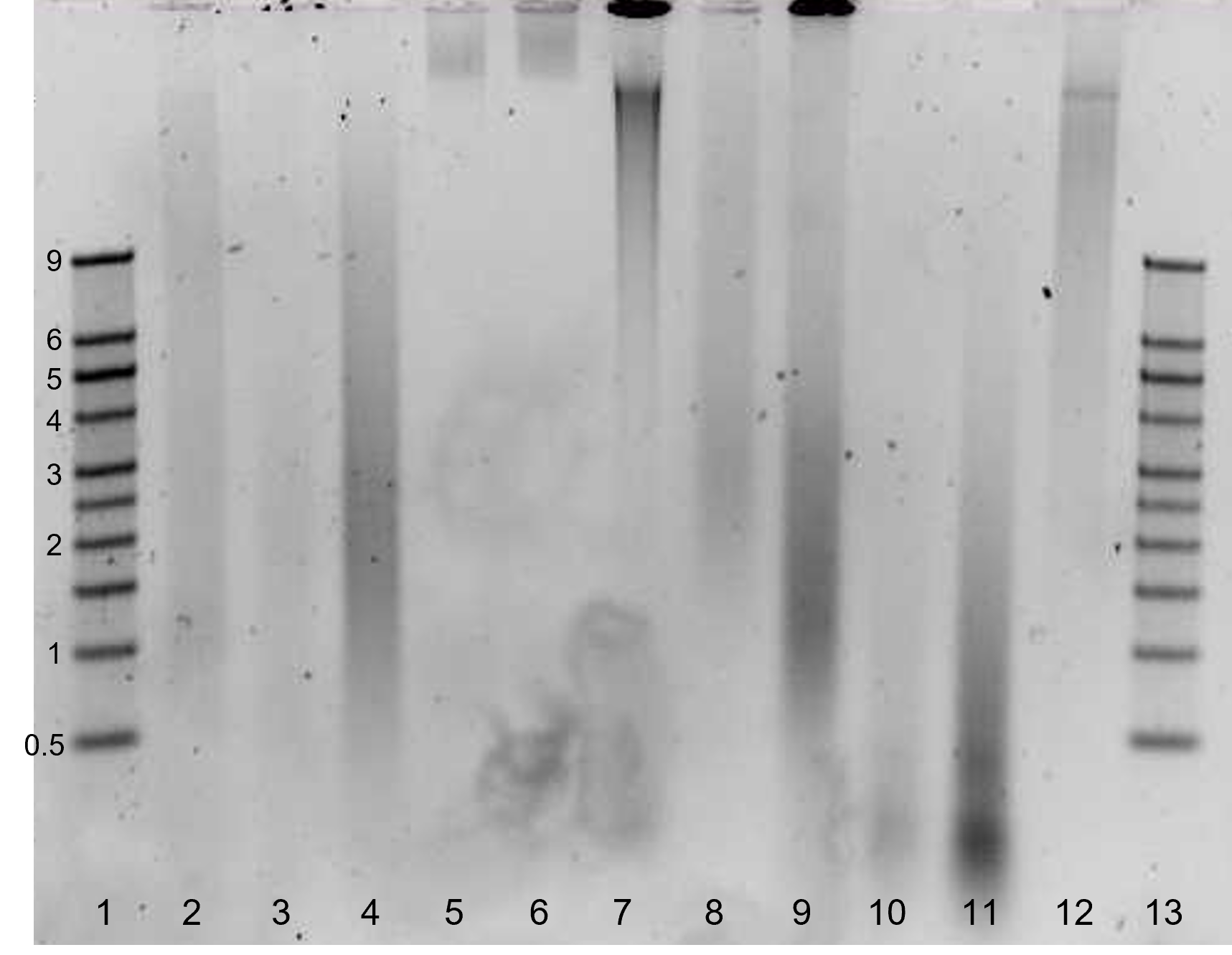
